# When feedback backfires: investigating neurofeedback effects in a closed-loop auditory attention decoding paradigm

**DOI:** 10.64898/2026.04.28.721343

**Authors:** Iustina Rotaru, Simon Geirnaert, Nicolas Heintz, Alexander Bertrand, Tom Francart

## Abstract

Selective auditory attention decoding (AAD) enables tracking which of multiple concurrent speakers a listener attends to and is a key building block for neuro-steered hearing devices. While AAD integrated in a closed-loop system with real-time neurofeedback (NFB) is hypothesized to improve decoding through neural adaptation and error-correction behaviour, the short-term behavioral and algorithmic impact of such a bilateral human-machine interaction remains poorly understood. Here we evaluated the effects of NFB on AAD accuracy and user experience in a single-session AAD paradigm with online NFB involving nineteen participants. They performed a selective listening task with enforced attention switches across four conditions: open-loop (OL), closed-loop with auditory gain feedback (CLA), closed-loop with visual feedback (CLV), and a condition with pseudo-auditory gain control (psCLA) decoupled from the participants’ individual neural activity. AAD was performed online using both subject-specific and subject-independent linear decoders on 5 s sliding windows, followed by Hidden Markov Model post-processing. Online analysis showed comparable decoding performance across all conditions. However, offline posthoc analysis using subject-independent decoders revealed that AAD accuracy in the CLA condition was significantly lower than in the OL baseline. Subjectively, participants reported that CLA was significantly more distracting and required higher switching effort. Crucially, a causal analysis of the psCLA condition found no robust evidence that higher audio gains inherently improve decoding accuracy. Our results demonstrate that within a single-session paradigm with rapidly varying feedback cues, auditory neurofeedback may degrade AAD performance by increasing cognitive load and distraction. These findings suggest that suboptimal feedback can impede rather than facilitate learning. We conclude that more accurate and stable decoders and longitudinal, multi-session training protocols are likely essential prerequisites for achieving beneficial neurofeedback effects in closed-loop auditory attention systems.

## 1 Introduction

In noisy “cocktail party” environments, the brain distinctly encodes sound sources based on attention. Specifically, the attended signals exhibit stronger neural representations than unattended ones^1,2^. This phenomenon enables Auditory Attention Decoding (AAD), which identifies the target, attended source from neural recordings like electroencephalography (EEG)^3,4^ or magnetoencephalography (MEG)^5^. The primary application of AAD is in neuro-steered hearing devices (nsHD), i.e., devices that dynamically amplify a user’s target speaker while suppressing other speakers and background noise^6^. Realizing such devices requires low-latency, closed-loop integration of AAD with real-time speech enhancement to provide immediate acoustic feedback based on the decoded attentional state^7,8^.

While AAD is advancing rapidly from linear methods^3,4,6^ to more complex deep learning architectures^9–11^, research remains largely focused on algorithmic robustness and cross-subject generalization^12,13^. However, translating these advancements into clinical nsHAs requires moving beyond offline analysis. Evaluation must also shift toward more ecological, out-of-the-lab settings characterized by frequent attention switches, a more complex listening environment with multiple distractors and reverberation, as well as miniaturized, mobile EEG systems^14–18^. Crucially, these systems must be validated with hearing-impaired individuals, the primary end-users, to account for altered neural speech processing^10,19,20^.

A critical, yet understudied dimension of AAD is the bilateral human-machine interaction in closed-loop systems, where real-time audio or visual feedback is provided based on decoded attention^21^. It is generally expected that such a closed-loop interaction could trigger neurofeedback (NFB) effects: by continuously interacting with the system and observing the feedback, the users learn how to regulate their brain activity in order to maximize an explicit target outcome, e.g., the level of the attended sound source and the overall decoding accuracy. We hypothesize that these learning effects may occur on different timescales: (1) on short-term, subjects can adopt an error-compensation strategy whenever they notice a unexpected mistake in the decoding algorithm and (2) on long-term, learning can manifest through the development of adaptive cognitive strategies over multiple sessions to improve system control. While both timescales contribute to what we define as the ‘neurofeedback effect’ (the iterative improvement of decoding through neural adaptation), the present study focuses exclusively on the former, short-term learning strategies.

Despite its potential, closed-loop AAD literature remains scarce mainly due to the technical complexity of the real-time implementation and validation. Existing studies involving online AAD with neurofeedback vary significantly in methodology and research objectives. Zink et al.^22^ showcased the earliest proof-of-concept for online AAD neurofeedback with mobile EEG and visual-feedback only, utilizing a longitudinal design (spanning four weeks) to test the potential training effects in ecological settings (the participants’ homes). They found early evidence that training might improve AAD performance, though their sample size was small (only two subjects) and it was difficult to distinguish true neural self-regulation from increased motivation or effort. Aroudi et al.^7^ realized the first closed-loop, cognitively-driven gain controller that directly modulates the gain levels of presented speakers based on probabilistic AAD. They systematically compared open- and closed-loop decoding and signal-to-interference (SIR) improvement in an attention-switching AAD paradigm. Although they demonstrated the system’s technical feasibility, they found no significant difference in decoding performance, subjective listening effort and speech understanding between open-loop and closed-loop sessions, thus indicating no clear behavioral benefit of neurofeedback. However, they show that the closed-loop system objectively improves the SIR between the attended and the unattended speaker, yet with of a significant delay in detecting attention switches (ranging between 8-36 s). Hjortkjær et al.^8^ integrated EEG decoding and microphone-based beamforming into a near-real-time neuro-steered hearing-instrument prototype, evaluating attention-controlled gain steering in normal-hearing and hearing-impaired listeners with audio-only and audiovisual speech. While demonstrating that real-time neuro-steered gain control is technically viable, they found that the audiovisual stimuli did not significantly improve the decoding performance w.r.t. the audio-only stimuli. They also highlighted challenges w.r.t. subject variability, the cost of misclassifications, speed–accuracy trade-offs and need for better decoding and user interfacing. Finally, Haro et al.^23^ used a closed-loop AAD-driven paradigm with gain control as a neurofeedback training tool aimed to understand the neural basis of attention improvement. In a single-session NFB training protocol, they manipulated the unattended talker’s gain and showed that closed-loop training selectively suppresses the neural tracking of the unattended speech and therefore reduces its decoding accuracy (p = 0.02, Cohen’s *d* = -1.29 for the unattended speaker correlation metric and p = 0.01, Cohen’s *d* = -1.56 for decoding accuracy of the unattended speaker). At the same time, they found no evidence of enhanced tracking (either in the correlation metric or decoding accuracy) of the attended speech between the first and second half of the experimental session. They however do not perform a behavioral assessment of the closed-loop system.

Expanding upon previous AAD-NFB research, we specifically aim to evaluate the short-term error-compensation effect mediated by neurofeedback (NFB) in a single-session AAD paradigm with two competing speakers and within-trial attention switches. To this end, we implemented a closed-loop AAD system that provides instantaneous feedback via neurally-driven gain control. To probe for NFB effects elicited by different feedback modalities, we propose a tailored protocol that includes a baseline, open-loop condition (OL, no feedback) and three closed-loop (CL) conditions: (1) with audio feedback (CLA), where the gain of the attended speaker increases relative to the unattended speaker, (2) with visual feedback (CLV), where a horizontal slider continuously moves towards the side of the attended speaker, and (3) with acoustic pseudo-feedback (psCLA). In the latter, the presented feedback is also implemented as a change in the relative gain between the two speakers, but, in contrast to CLA, the feedback is fully decoupled from the participants’ neural responses. Instead, it is derived from neural data recorded in the separate OL trials, whose left-right attention and switching patterns match those in psCLA. This approach preserves the realism and ecological validity of the feedback, while participants remain unaware of this manipulation, as the audio gains still generally match the underlying attention pattern of the participant. Hence, psCLA can be regarded as a ‘realistic sham’ condition.

Our main hypothesis is that CLA and CLV trials will yield significantly higher decoding accuracies than OL and psCLA trials, due to possibly two different underlying mechanisms. Firstly, we suspect that in CLA, a positive reinforcement effect may occur, as the amplification of the target source might lead to improved focus and less distraction to the tuned-down sources, thereby enhancing the neural tracking and the subjective listening experience. Secondly, we hypothesize that *active* error compensation will occur in the CLA and CLV trials. Since no AAD algorithm is error-free, either due to inherent decoder’s errors or momentary distraction on the subjects’ part, immediate feedback would allow subjects to detect misclassifications or moments of distraction in real-time. We thus expect users to adopt reactive mental strategies such as increasing cognitive effort or intensifying focus to “correct” the decoder’s output. Given these two mechanisms, we overall predict that the presence of correlated feedback (in CLA/CLV) will result in superior AAD performance compared to the absence of feedback (OL) or the non-contingent feedback from the ‘sham’ feedback condition (psCLA).

Beyond objective decoding metrics, we aim to also qualitatively evaluate the impact of neurofeedback on subjective markers of speech perception in noise. Specifically, we assess speech intelligibility, listening and switching effort, degree of distraction, as well as the user’s sense of agency (perceived control) while operating the CL system. Hereby we hypothesize that in CLA, the subjects will report significantly higher speech intelligibility, reduced listening effort and degree of distraction compared to all the other conditions, since the auditory feedback is explicitly designed to improve the signal-to-noise ratio (SNR). Additionally, we hypothesize that the objective and subjective metrics will be positively correlated, i.e., that higher decoding accuracies will correspond to an increased subjective sense of system control and accuracy estimation (subject-predicted performance).

Finally, we aim to address a critical confound: whether the hypothesized neurofeedback effect (higher accuracy in CLA) is a by-product of enhanced acoustic gains or a result of neural adaptation. While the system’s design ensures that better decoding naturally leads to improved gains, the inverse relationship - whether higher gains causally drive better decoding - remains to be elucidated. By analyzing the causal relationship between gain levels and AAD accuracy, we seek to disentangle the purely acoustic benefits of the system from the cognitive-neural gains induced by the closed-loop interaction.

## 2 Methods

### 2.1 Participants

A total of 31 participants were recruited for this study. The selection criteria were: a normal hearing profile (hearing thresholds were assessed using pure tone audiometry and confirmed to be ≤ 25 dB HL at all octave frequencies from 125 to 8000 Hz), no attention deficits and Flemish as mother tongue. Twelve participants were not included in the final analysis since they took part in initial pilot tests to determine the right test protocol, conditions and parameters, such as audio presentation levels, hyperparameters of the gain control system etc. The final analysis was thus performed on the data from nineteen participants (9 males, 10 females), which all took part in the same protocol with the same hyperparameters. Their age ranged between 19-65.9 years (with mean age of 31.04 ± 14.79 years). The experiment was conducted in accordance with the principles embodied in the Declaration of Helsinki and was approved by the Ethics Committee Research UZ/KU Leuven, Belgium (project reference S57102). All participants signed a written informed consent for voluntarily participating in the study.

### 2.2 Experimental protocol and setup

Initially, the participants were given a short presentation about the scope and tasks of the NFB experiment, whose backbone is the two competing-speaker AAD paradigm: the main task is to listen to the target speaker (marked in green) and ignore the other one (marked in black). The subjects were informed that a brain decoder will be used to provide real-time feedback based on the predicted auditory attention (in the closed-loop setting), and that they have to find a good mental strategy to control either the visuals or the audio gains to enhance the target speaker. They were asked to minimize frowns, chewing, face, head and body movements to minimize the artefacts in the EEG data.

Additionally, to facilitate participants’ constant engagement with the system, a second task they had to accomplish was to apply a mental strategy to maximize the decoding accuracy score, which was displayed on-screen at the end of each trial. They were informed that the accuracy score measures the percentage of time when the system correctly decoded their auditory attention to the green speaker during that trial and that the higher this score, the better they were able to control the system with their brain. Just before displaying the genuine accuracy score for each trial, they were asked to predict this value with a number between 0-100 % based on their perceived performance (as reflected by the feedback cues) in that trial.

The general experimental timeline is illustrated in fig. 1. The neurofeedback (NFB) experiment was divided in two main blocks of 4 trials each. Each trial lasted for 8 min and was assigned a certain condition type, i.e. based on the presented feedback modality. In the experimental design, we included four condition types: open-loop without feedback (OL), closed-loop with visual feedback (CLV), closed-loop with audio feedback (CLA) and closed-loop with audio pseudo-feedback (psCLA). The order of these conditions within a block was pseudo-randomized, with the only constraint that the OL trial always preceded the psCLA trial (for reasons outlined below). The subjects were explicitly informed about the feedback modality they will receive (audio, visual or no feedback) at the start of each trial.

**Figure 1.**
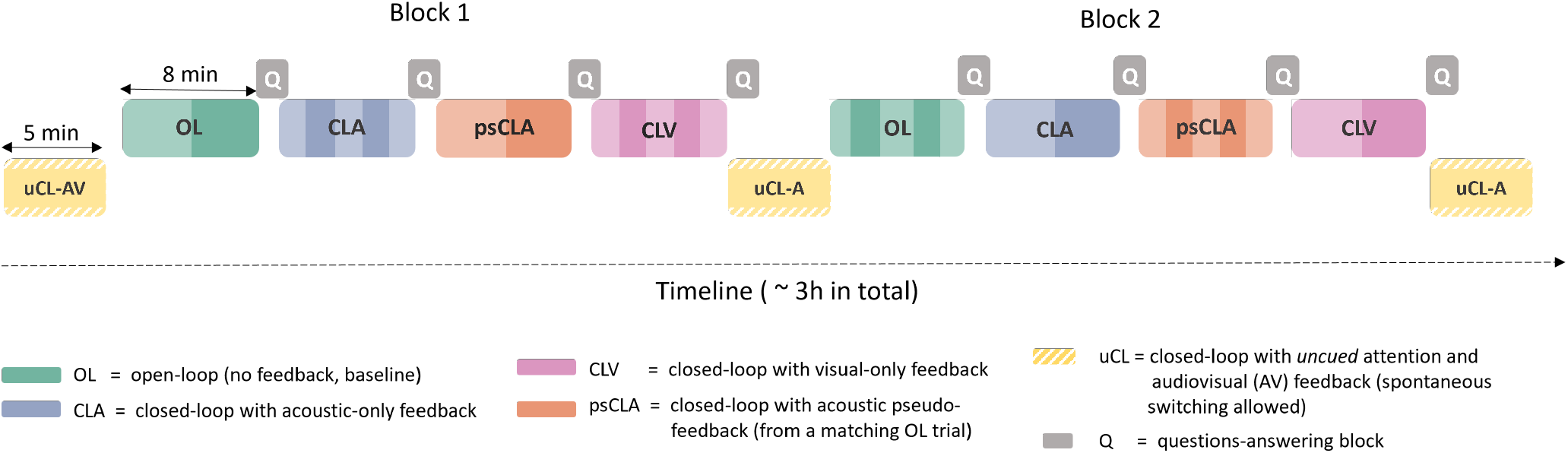
Experimental protocol with an exemplary sequence of trials. Each color denotes a different condition type, as marked in the legend. The shaded areas within each trial delimit the periods with sustained attention to one of the speakers, in between the switching moments. 1-switch and 3-switch trials were always alternatively presented. The order of the conditions within each block was pseudo-randomized for each subject, ensuring that the OL trial always preceded the psCLA trial.

The OL condition served as the baseline - there was no feedback or interaction with the system, and the subjects’ only task was to actively listen to the green speaker and ignore the other.

In CLV, a visual green slider was presented, which moved step-wise along a virtual horizontal line adjoining the two L-R speaker icons presented on-screen (see fig. 2, fig. 3). The slider’s exact position within the horizontal line was determined based on the decoder’s output (details in section 2.6). For example, if the L-speaker was decoded over several consecutive windows, the slider would move consistently towards the icon of the L-speaker with a step-size proportional to the decoder’s output. Whenever a switch in attention was detected by the decoder, the movement direction of the slider would change towards the newly attended speaker. In this condition, the participants’ task was two-folded: to actively listen to the target speaker marked in green, and to control and keep the moving slider consistently towards the side of the green target speaker.

**Figure 2.**
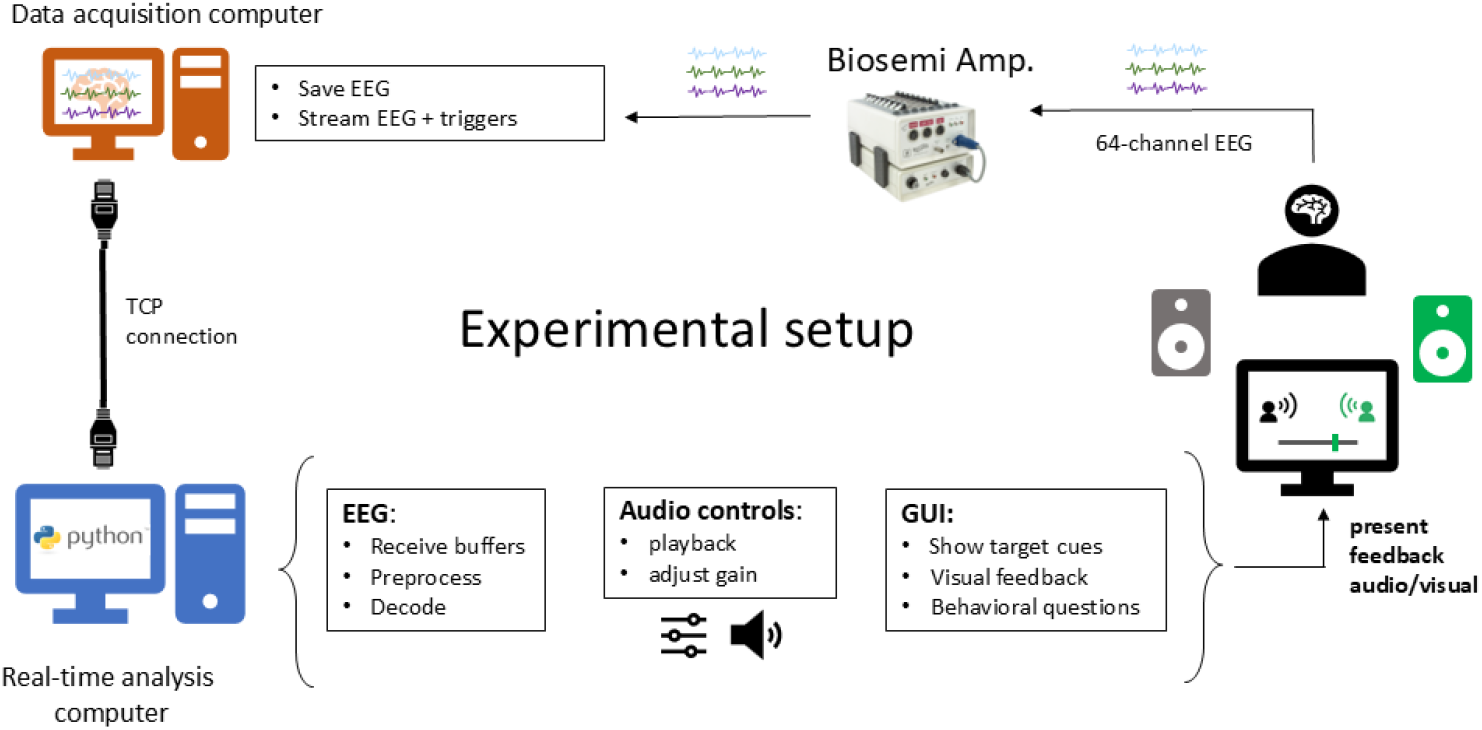
Experimental setup of the closed-loop AAD system with real-time visual/audio feedback

**Figure 3.**
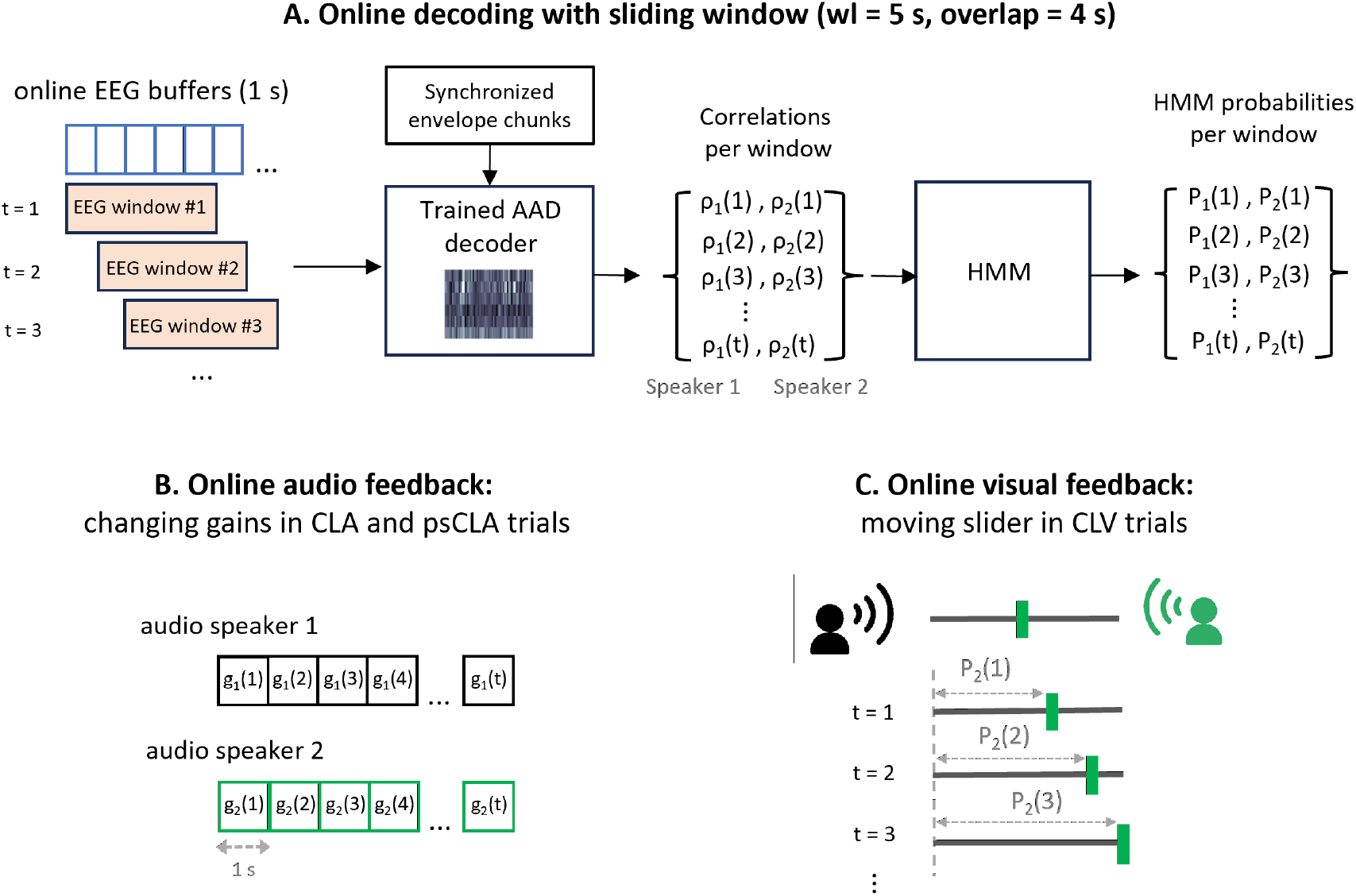
Online decoding pipeline and feedback presentation. **(a)**: overview of the online AAD decoding with a sliding window (decoding window length = 5 s, overlap = 4 s, stride = 1 s) and the post-processing of AAD correlations with an HMM. The HMM output probabilities are used to determine the audio feedback in the form of changing gains **(b)**, and the visual feedback in the form of a visual moving slider **(c)**.

In the CLA condition, the presented audio feedback consisted in changing the gains of the two speakers in a step-wise manner, such that the intensity of the attended speaker was enhanced and the intensity of the unattended speaker was reduced (the gain change system is presented in more detail in section 2.6). The participants’ task again was two-folded: to actively listen to the green speaker and to control the gain change system with their brain such that the volume of the green speaker is maximized.

Finally, the psCLA condition was ‘disguised’ to the participants and presented identically as the CLA condition (in terms of task, instructions and feedback), but unbeknownst to the participant, they did not control the gain of the two speakers themselves. Instead, we used a pre-computed gain trajectory based on the EEG data recorded in the OL trial from the same block and applied these gains to the speakers in the psCLA trial. Notably, even though we used different stimuli in the 2 trials, we specifically enforce the attention switching patterns between the OL and psCLA trials to be well-matched (i.e., these trials have the same sequence of attended speakers and number of switches, e.g., L-R-L-R), such that the feedback sounds very realistic and thus gives the illusion of control. We introduced the psCLA as a ‘control’ condition in order to disentangle potential effects on AAD accuracy due to mere feedback presentation (even if uncorrelated from behaviour) vs. due to genuine feedback (CLA and CLV), which is contingent upon the participant’s real-time decoded behaviour.

To provide initial familiarization with the system as well as more training opportunities during the NFB session, three shorter closed-loop trials with uncued attention (uCL) were included at the start and end of the NFB session, as well as in-between the two main experimental blocks. In these trials, the subjects were free to attend to whichever speaker they wanted and to switch attention in between the 2 speakers, with the only constraint that the switching should not happen too fast, i.e. they should leave at least 10 seconds in between two consecutive switching moments. These uCL trials were not further used in the main analysis due to the lack of ground-truth attention labels. However they served as a reference in a final questionnaire, in which subjects were asked to rate their overall experience, indicate systems’ strengths, limitations, aspects of further improvement and which mental strategies they used to control the system.

To make the audio scenario more realistic and probe how the system reacts in different scenarios with various degrees of attention switching, we chose to have two trials for each condition, with either one or three switches in attention. In the 1-switch trials, the attention was switched between the speakers once at mid-trial, i.e., at 4 min. In the 3-switch trials, the attention was switched every 2 min. In the experimental design, we ensured that the number of attention switches always alternated between two consecutive trials (see fig. 1). The switches were visually cued by presenting a green arrow that changed direction towards the new to-be-attended speaker.

After every trial, participants answered questions about the content of the target attended speaker (to ensure compliance with the attention task), and were asked to perform a self-assessment of their speech intelligibility, listening effort, switching effort, degree of distraction and sense of control of the system by rating each of these subjective factors with a number between 1-10. An English translation of these subjective questions is presented in the Supplementary Material. Short breaks of 2-3 minutes were taken after every trial and a larger break of about 10 min was offered mid-experiment to promote a reset of the subjects’ attention span and motivation. In summary, the entire experiment included the introduction, NFB session, breaks and question-answering blocks and lasted for about 3h in total.

### 2.3 Stimuli

We used podcasts from ‘Universiteit van Vlaanderen’ as audio stimuli^24^. For each trial, we randomly selected two different stimuli from a pool of downloaded podcasts, ensuring that both the speakers and the attended/ignored content were always new. We only considered podcasts with male voices to avoid mixed-speaker’s sex effects on attention^25^. Each stimulus was presented from a separate loudspeaker via a Behringer soundcard (U-PHORIA UMC1820, Germany). The two loudspeakers were positioned at 1.4 m in front and ±14° azimuth angle relative to the seated participant.

In the preprocessing stage, we trimmed all silences to 200 ms in order to avoid periods of longer silences when the subjects could be more easily distracted by the other speaker. The stimuli were further normalized to -27 dB SPL and were finally amplified with predetermined calibration gains to reach the desired output intensity level.

The stimuli were presented at different intensities depending on the trial: in both OL and CLV conditions, the two stimuli were presented with an equal intensity level of 57 dBA. This was considered the baseline intensity, and was chosen because it was a comfortable sound level in preliminary pilot tests. For the CLA, psCLA and uCL trials, the acoustic gains applied to both speakers changed every second, depending on the decoder’s output (details in section 2.6). Altogether, the gains ranged between [-6, +6] dB, reaching thus sound intensities between [51, 63] dBA, which were empirically determined as minimum and maximum amplification levels that are comfortable and that render audible speakers to which it is still possible to switch attention.

### 2.4 Online EEG acquisition and preprocessing

Our experimental setup is illustrated in fig. 2. 64-channel EEG data were recorded at 1024 Hz with a Biosemi amplifier (The Netherlands) and the Actiview software on our data acquisition computer. The data buffers were further transmitted in real-time via a TCP connection to our analysis computer, which ran Python scripts that handled the real-time EEG analysis, audioplayer controls and the graphical user interface as simultaneous processes in a multiprocessing coding framework. The audio stimuli and a pulse trigger signal were played simultaneously via a Behringer UMC 1820 soundcard. The triggers synchronized the EEG data with the audio by encoding a timestamp into the EEG stream at one-second intervals throughout stimulus playback.

The incoming EEG data was preprocessed in buffers of 1 s and decoded with a sliding window of 5 s and a stride of 1 sec. Thus, every new EEG buffer was preprocessed, then concatenated with the previous 4 EEG buffers, and finally submitted to the AAD decoder functions. This enabled us to apply our decoder and to change the feedback signal (slider position or acoustic gain) every 1 s, such that the system felt responsive and interactive to the users. Note that due to our sliding window approach with 4 s overlap, the consecutive decoding windows aren’t completely independent.

The sequential preprocessing steps per EEG buffer were: downsampling to 64 Hz, re-referencing to the common-average of all channels and bandpass-filtering in the 1-9 Hz range. The downsampling was performed after applying a lowpass antialiasing filter, namely a Chebyshev type I filter of order 8 and a higher cutoff frequency of 25.6 Hz. The bandpass filter was a digital IIR Butterworth filter of order 4 and a passband of 1-9 Hz. Both filters were defined by second-order-section (“sos”) coefficients and their filter states were updated buffer-by-buffer in order to avoid signal discontinuities and edge artefacts.

### 2.5 Auditory Attention Decoding (AAD)

In this study, we used linear decoders based on the stimulus reconstruction paradigm^3,6^. Esentially, the goal is to reconstruct the speech envelope of the attended speaker from the neural signals. The spatio-temporal decoder 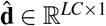 is found by minimizing the squared error between the attended speech envelope **s** ∈ ℝ^*T* ×1^ and the reconstructed envelope from time-lagged versions of the EEG signal **X** = [**X**_1_ … **X**_*C*_] ∈ ℝ^*T* ×*LC*^ (where *C* is the number of EEG channels, *L* the number of timelags and *T* the number of time samples in the training data):

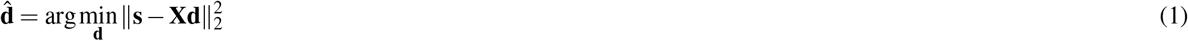

with

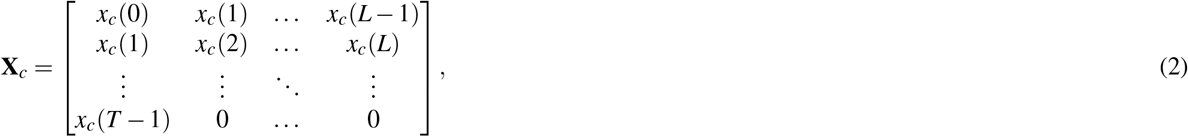

the EEG matrix that stacks the EEG signal at channel *c* over *T* samples and *L* timelags. The timelags are added to account for the neural processing delay of an auditory stimulus, whereby the stimulus always predates the neural response. Here we used positive timelags of up to 250 ms (corresp. to 16 samples for a sampling rate of 64 Hz), which is a typical choice in the literature^6^. The analytical least-squares solution of eq. 1 is:

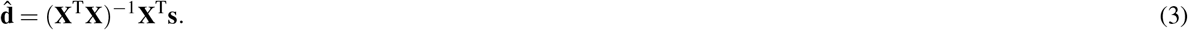

To check if the type of decoder impacts the accuracy scores in the different NFB conditions, both generic, subject-independent (SI) decoders, as well as subject-specific (SS) decoders were used online on a different subset of subjects. Although SS decoders are generally more accurate^6^, SI decoders remain highly relevant for practical applications due to their ability to eliminate or minimize the calibration phase. Thus, the participants were split in two experimental groups, depending on the type of AAD decoder used to give online feedback: the *SI group* (n=10), which received online feedback generated by the pre-trained SI decoder, and the *SS group* (n=11), which received online feedback with an ad-hoc trained SS decoder. Notably, two participants were tested twice on two separate days/sessions - once with the SS and once with the SI decoder. Per subject and session, the assigned decoder remained static - no online adaptation or re-training occurred during the trials.

The SI decoder was trained on the AAD-KUL dataset^26^, which was recorded with the same EEG system. Conversely, for training the SS decoder for the SS subjects group, we recorded EEG data in four additional calibration (CAL) trials of 5-min each (which preceded the main NFB experiment), totalling 20 min of training data. There were no attention switches in these calibration trials, but attention was overall balanced between the L & R speakers across trials. Notably, the CAL trials were recorded in the same conditions as the OL trials from the main experiment, i.e., no feedback was presented.

Per trial and subject, the pre-trained decoder (either SS or SI) was then applied on-the-fly on every new decoding window of 5 s (with a stride of 1 s and an overlap of 4 s). The output of the decoder was a vector of two correlation values that marked the strength of the linear mapping between the reconstructed envelope and each of the two speakers’ envelope. In each test window, we selected as attended speaker the one which yielded the highest correlation value.

For a comprehensive retrospective analysis, we employed two different SI decoders to evaluate offline performance across various non-overlapping window lengths. This offline validation aimed to assess the generalizability and robustness of our AAD system under a more standardized test setting, specifically eliminating the temporal dependencies introduced by the overlapping windows used in the online implementation. Additionally, we aimed to re-evaluate all subjects with a uniform benchmark decoder for comparative consistency. The first offline decoder was trained on the AAD-KUL dataset^26^, i.e., it was the same decoder used in the online sessions of the SI group, but was applied retrospectively to the full cohort. The second offline decoder was trained posthoc using a leave-one-subject-out cross-validation scheme on the OL and CAL trials from the current study’s cohort (hence, this is the ‘study-specific’ SI decoder). By restricting the training set only to the CAL and OL trials, we ensured the decoder remained unaffected by potential neurofeedback (NFB) effects or adaptive strategies developed by subjects during the feedback sessions. A detailed summary of the training and testing configurations for all decoders used in this study is provided in Table 1.

**Table 1.**
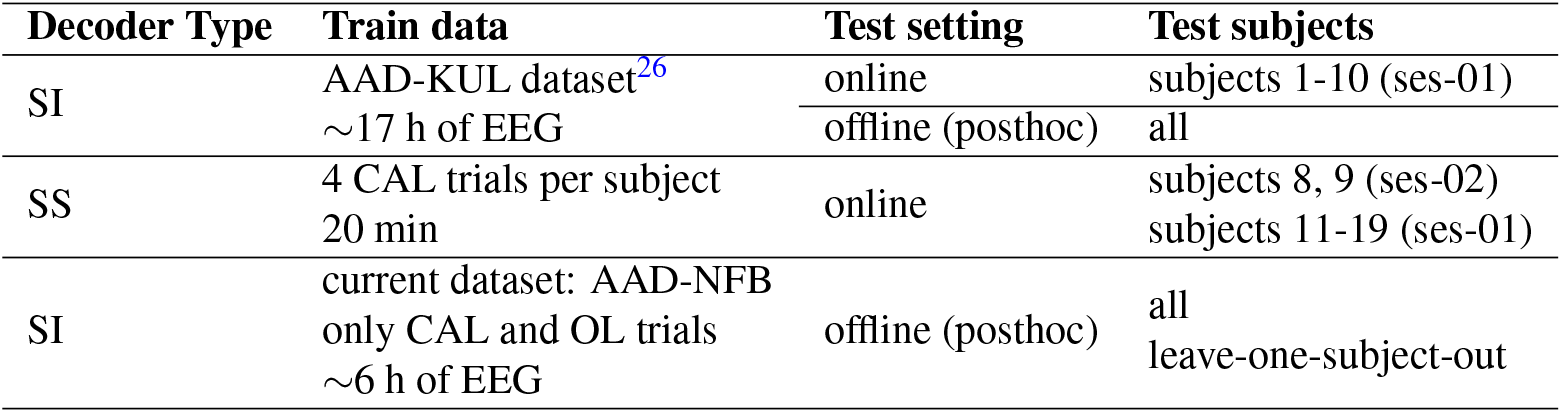
Overview of all AAD decoders used in this study. SI = subject-independent decoder, SS = subject-specific decoder. CAL = calibration, OL = open-loop.

#### Envelope calculation

The stimuli envelopes were pre-computed and loaded into the EEG analysis script in the beginning of each trial. Initially, the audio stimuli were downsampled to 16 kHz and passed through a gammatone filterbank, which roughly approximates the spectral decomposition as performed by the human auditory system^27^. Per subband, the audio envelopes were extracted by taking the absolute value of the stimulus waveform (full-wave rectification) and compressing its dynamic range with a power-law operation with exponent 0.6 (as proposed in^27^). The resulting subband signals were then summed to construct a single broadband envelope, which was further downsampled to 1024 Hz to match the sampling rate of the online streamed EEG. Finally, the envelopes were split in smaller buffer windows of 1 s and submitted to the same sliding-window preprocessing procedure as the EEG data (see the EEG preprocessing steps from section 2.4). The EEG and envelopes were synchronized in the online setup based on triggers received every second.

### 2.6 Smooth audio and visual feedback with HMM post-processing

We rely on the computed correlations in every newly decoded window to present feedback, either in the form of audio gains for the attended and unattended speaker (in CLA trials), or as a change in the visual slider position (in CLV trials). However, instead of using the raw correlation values computed within each 5 s window, which are inherently noisy, we post-process them with a real-time, causal Hidden Markov Model (HMM), as proposed by Heintz et al.^28^. This model aims to improve the accuracy of instantaneous AAD predictions by modeling the temporal structure of attention, based on the principle that a listener is much more likely to sustain attention to the same speaker than to frequently switch attention. We note that the 4 s overlap between consecutive 5 s windows violates the temporal independence condition within the HMM algorithm’s theoretical foundation, yet we empirically observed that the HMM still gave good results that were better compared to a setting with either 1 s or 5 s non-overlapping windows.

In brief, the HMM operates on the sequence of AAD scores, in our case correlations, *ρ*_*j*,0:*t*_ = [*ρ*_*j*,1_ … *ρ*_*j,t*_], where *ρ*_*j,t*_ is the correlation value for speaker *j* in the decoding window *t*. The goal of the causal inference is to compute the conditional probability *P*_*j*_(*Ŝ*(*t*) = *S*_*j*_|*ρ*_*j*,0:*t*_), which is the probability that the listener is attending to speaker *S*_*j*_ in window *t*, given all observed correlations to speaker *S*_*j*_ up to time *t*. This provides a smoothing effect on the AAD decisions. This probability is computed recursively and causally using the Forward algorithm^28^, which calculates the joint probability *α*_*j*_(*t*):

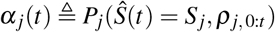

The recursive calculation of *α*_*j*_(*t*) is given by:

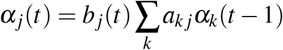

where:

- *a*_*kj*_ ≜ *P*_*j*_(*Ŝ*(*t*) = *S*_*j*_ *Ŝ*(*t* 1) = *S*_*k*_) is the transition probability, defined by the underlying Markov chain structure of attention. This probability reflects the likelihood of switching from speaker *S*_*k*_ to speaker *S*_*j*_. For a simple Markov chain with 2 speakers, *a*_*kj*_ is typically defined by a small switching probability *a*_12_ = *a*_21_ = *p*_*switch*_.
- *b*_*j*_(*t*) ≜ *P*_*j*_(*ρ*_*j*_(*t*)|*Ŝ*(*t*) = *S*_*j*_) is the emission probability, which models the likelihood of observing the correlation *ρ*_*j*_(*t*) given that *S*_*j*_ is the attended speaker. This distribution is assumed to be Gaussian, whereby *µ* and std were empirically estimated from data of five pilot subjects in the current protocol.

The recursion starts at *t* = 0 with the initialization:

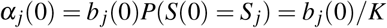

assuming all *K* speakers (*K* = 2) have an identical prior probability to be attended. Once *α*_*j*_(*t*) is computed, the final conditional probability that speaker *S*_*j*_ is attended in window *t* is calculated by normalizing *α*_*j*_(*t*) over all possible speakers *K*:

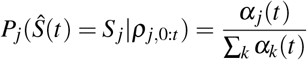

For practical implementation, especially with longer observation sequences, the algorithm is performed in the logarithm domain (using log probabilities) to ensure numerical stability. The causal HMM algorithm is equivalent to skipping the backward pass in the forward-backward algorithm (setting *β*_log, *j*_(*t*) = 0 in Algorithm 1 from Heintz et al.^28^).

*p*_*switch*_ is a tunable hyperparameter that controls how fast the HMM will detect an attention switch. The lower this value, the more stable the resulting gain trajectories will be, however at the cost of a longer detection time in case of a true switch in attention. To choose an optimal value for *p*_*switch*_, we conducted an offline posthoc analysis using EEG data from the first five pilot participants. We tested values between (10^−2^, 10^−5^) in logarithmic steps: for each value, we computed posthoc gain trajectories per trial and evaluated them by computing the stability and switch-detection time. Stability was defined as the percentage of windows within a trial in which the gain of the attended speaker was at or above the comfort level (here set to 80% of the maximum allowed sound level). Switch detection time was defined as the average time between when the listener switched their attention and when the HMM correctly predicted the newly attended speaker (i.e., assigned it the highest probability). Based on these analyses, we selected *p*_*switch*_ = 0.01 for the online NFB experiment, as it provided a favorable balance between stability and switch detection speed, which was also visually noticeable in the plotted gain trajectories.

The online decoding with the sliding window and the resulting feedback presentation is illustrated in fig. 3. In the CLA trials, the HMM was deployed online and used to determine the audio gains presented in the next 1 s buffer based on the correlation values in the present decoding window. In the case of *K* = 2 speakers, the HMM yields two sequences of probabilities, *P*_1_ and *P*_2_, one for each speaker. To obtain the actual gain values to apply to each speaker in the next audio buffer, we linearly mapped these probabilities to our desired gain range of [-6, +6] dB by means of linear interpolation, obtaining the sequence of gains *g*_1_ and *g*_2_ for speakers 1 and 2, respectively (fig. 3 B). An actual example of gain trajectories obtained with HMM is shown in fig 4. Note that the gains of the two speakers in each window *t* are always complementary, e.g. -5 dB and +5 dB. This is because for each window *t*, the HMM probabilities add to 1: *P*_1_(*t*) + *P*_2_(*t*) = 1.

**Figure 4.**
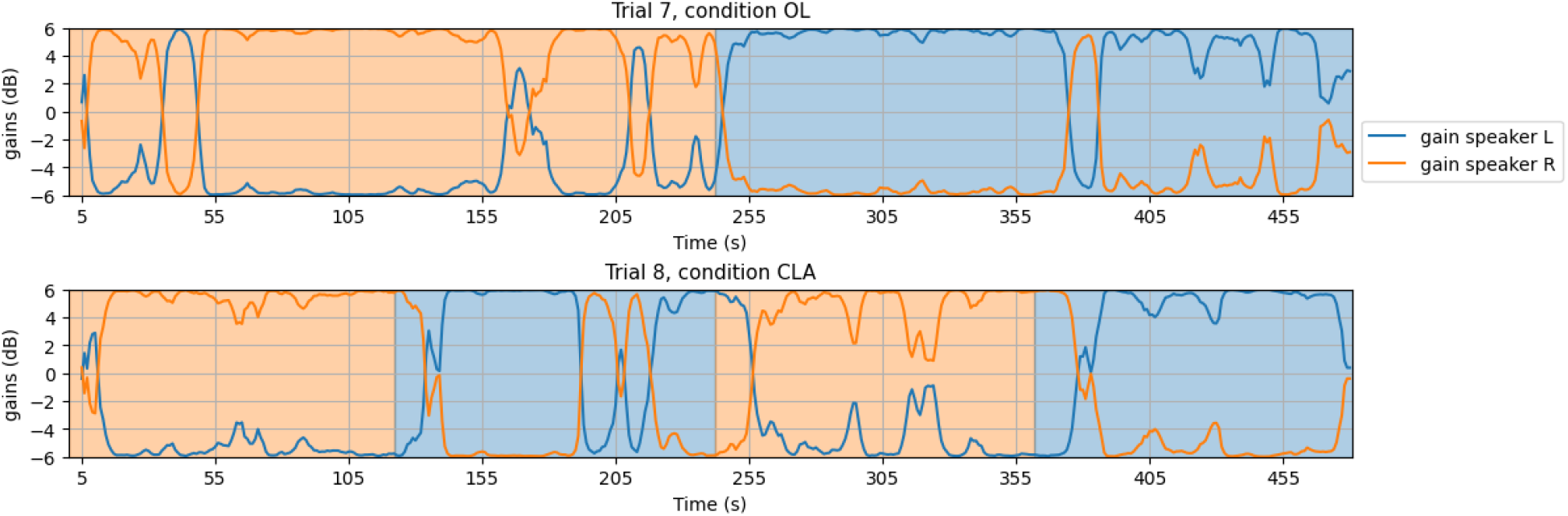
Illustrative gain trajectories for a CLA trial (online) and OL trial (posthoc) from a representative participant. The gains are relative to the baseline sound intensity level of 57 dB A. The background colour denotes the attention ground-truth, i.e. the L/R target speaker to which the subjects were instructed to pay attention (orange for the R speaker, blue for the L speaker).

In the psCLA trials, the presented gains are precomputed with an HMM applied offline on data from a matched OL trial. The resulting gain trajectory is pre-loaded at the start of the psCLA trial and for each new second of the stimuli playback, the corresponding gains from the gain trajectory are extracted and applied. For the subject, this mimicks the online decoding as performed in CLA, but behind the scenes, no genuine online decoding is in fact performed.

In the CLV trials, an HMM is also deployed online to provide visual feedback. However, instead of mapping the HMM output probabilities to gain values, this time, the output probability for the R-speaker determines the offset position of the visual slider, relative to the left slider-bar margin (as depicted in fig. 3 C). For consecutive decoded windows, the slider thus appears as moving step-wise across the horizontal bar, depending on which speaker is determined to be attended with the highest probability (as computed by the HMM algorithm).

Table 2 summarizes the feedback presentation per condition.

**Table 2.** Overview of the feedback presentation per condition.

| Condition | Presented feedback | Description |
| --- | --- | --- |
| OL | - | classic 2 speaker AAD paradigm (no feedback) |
| CLA | audio | genuine NFB with audio gain control based on HMM output probabilities $(P_1(t), P_2(t)) \rightarrow (g_1(t), g_2(t))$ |
| psCLA | audio | mimicks CLA, but no real NFB - the presented audio gains are pre-computed with HMM based on a matched OL trial |
| CLV | visual | moving slider - its position is determined by the R-speaker probability of the HMM: $P_2(t)$ |

### 2.7 Evaluation and statistical analysis

In the online setting, we computed three evaluation metrics per condition: the AAD accuracy (before HMM postprocessing), the HMM accuracy and the mean Signal-to-Noise ratio (SNR). The AAD accuracy is the percentage of correctly classified decision windows and is determined by comparing the window-wise decoder predictions with the ground truth attention labels. The HMM accuracy is computed as the percentage of windows where the HMM output probability of the attended speaker is larger than that of the unattended speaker (note that for the CLA trials, the HMM accuracy also corresponds to the percentage of decoded windows in which the attended speaker is louder than the unattended speaker). Significance levels for these accuracies were determined using the inverse binomial distribution (*α* = 0.05), taking into account the total amount of available test windows and a significance threshold of *α* = 0.05. Finally, the mean SNR is computed as the average gain difference (in dB) between the attended and unattended speakers across all decoded windows.

To evaluate the effects of the experimental conditions and decoder types, we employed Linear Mixed-Effects (LME) models. This approach was chosen to account for the hierarchical structure of our data, treating the multiple observations per participant as nested within each subject. For each outcome measure, we defined an LME model with *Condition* (OL, CAL, psCLA, CLV) and *Online Decoder Type* (SI vs. SS) as fixed effects, while Subject ID was included as a random intercept to account for inter-subject variability. The outcome metrics were the following scores averaged per condition across trials: AAD accuracy, HMM accuracy, mean SNR, as well as the behavioral scores (speech intelligibility, listening effort, switching effort, degree of distraction, perceived sense of agency and subjects-estimated accuracy). The LME models were fitted in R using the formula:

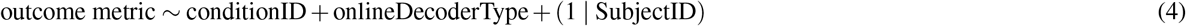

The significance of the fixed effects was assessed using a Type III Analysis of Variance (ANOVA) with Satterthwaite’s method for approximating degrees of freedom. Where significant main effects were identified, we performed posthoc pairwise comparisons using the *emmeans* package, applying Tukey adjustments for multiple comparisons.

### 2.8 Analysing the causal relationship between presented gains and AAD accuracy

Besides NFB effects influencing the AAD accuracy, we also hypothesize that higher gains for the attended speaker might inherently improve AAD performance, regardless of the subject’s neural modulation. Thus, to disentangle the NFB effects from gain change effects on AAD accuracy, we analysed the causal relationship between the presented gains and AAD accuracy, specifically looking at whether the current audio gain of the attended speaker leads to changes in the AAD accuracy in the current and future data windows.

To isolate this effect, we restricted this analysis to the psCLA condition only, because the gains are updated online, but they are decoupled from the subject’s current brain activity (as they are derived from EEG data of a different trial). We intentionally avoid the analysis of the CLA condition, as in CLA we cannot isolate a fully causal effect of gains on the AAD accuracy. This is because in CLA, the acoustic gains are updated online based on the correlation computed in the previous sliding windows, leading to an inherent mutual dependence of the gains and the AAD accuracy.

Thus, to conduct this posthoc analysis on psCLA trials, we used the same decoders that were used online per subject (either SS or SI decoders), but segmented the EEG data using *N* non-overlapping windows of length *WL*_*gain*_ ∈ {1, 5, 10, 20} seconds. For each window *n* ∈ {1,…, *N*}, we compute:

- 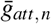: the average presented gain of the attended speaker within window *n*
- *ρ*_*att,n*_ and *ρ*_*unatt,n*_: the Pearson correlation coefficients for the attended and unattended speakers, respectively, computed offline with the SS/SI decoders

Subsequently, to probe the causal relationship between gains and AAD accuracy, we define two temporal mapping functions, ℳ_1_ (concurrent) and ℳ_2_ (forward):

1. Concurrent Mapping 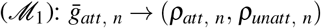
2. Forward Mapping 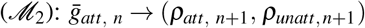

These mappings are used to pair gains with correlation outcomes. Mapping ℳ_1_ probes instantaneous effects of gain on AAD accuracy, whereas mapping ℳ_2_ essentially tests whether the influence of the presented gains extends beyond the current decoding window. A conceptual schematic overview of these two temporal mappings is presented in fig. 5.

**Figure 5.**
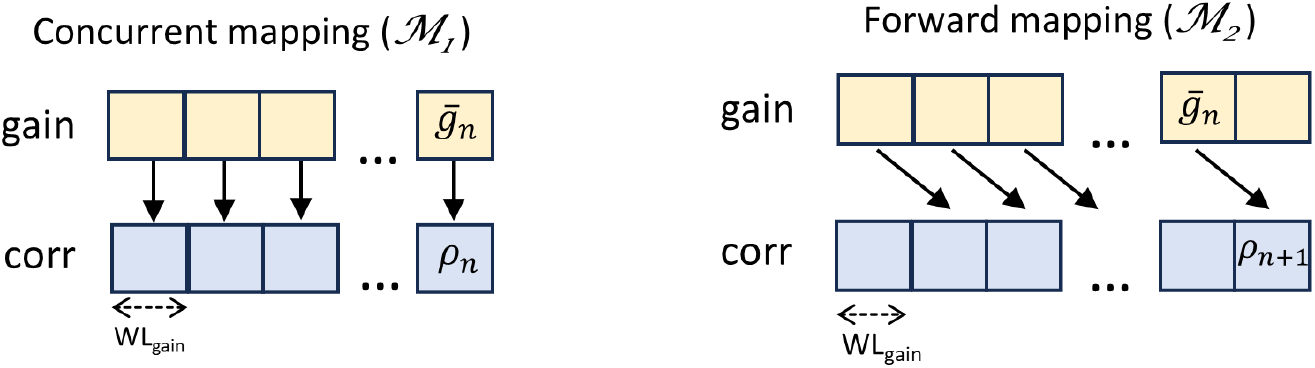
Schematic overview of the temporal mappings used to probe the causal influence of acoustic gains on the AAD accuracy in the psCLA trials

Per subject and window length *WL*_*gain*_, we then discretize the gain values 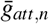 into a set of three bins *B* = {[−6, −2), [−2, 2), [2, 6]} dB, in order to capture effects from small, medium and high presented gains. For each mapping and gain bin *b* ∈ *B*, we pool together all triplets 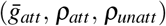 falling within the respective gain bin. Finally, the AAD accuracy for each bin is calculated as the percentage of windows where *ρ*_*att*_ *> ρ*_*unatt*_. We considered a small number of bins to ensure that enough datapoints are falling within each bin and thus obtain a good accuracy estimate per gain bin (per subject).

We performed this gain bin-accuracy analysis individually for each subject, and further aggregated the results per gain bin across subjects to capture inter-individual variability in the neural responses to the presented gain levels. For statistical comparison, Wilcoxon signed-rank tests are used to compare AAD accuracies across subjects between pairs of gain bins, with p-values adjusted for multiple comparisons using the Benjamini–Hochberg procedure.

## 3 Results

### 3.1 Online decoding: consistent performance in CL and OL conditions

Figure 6 illustrates the online performance metrics. To ensure a fair comparison, performance for the OL and psCLA conditions was evaluated posthoc using the same sliding-window and HMM hyperparameters used during the online CLA/CLV trials.

**Figure 6.**
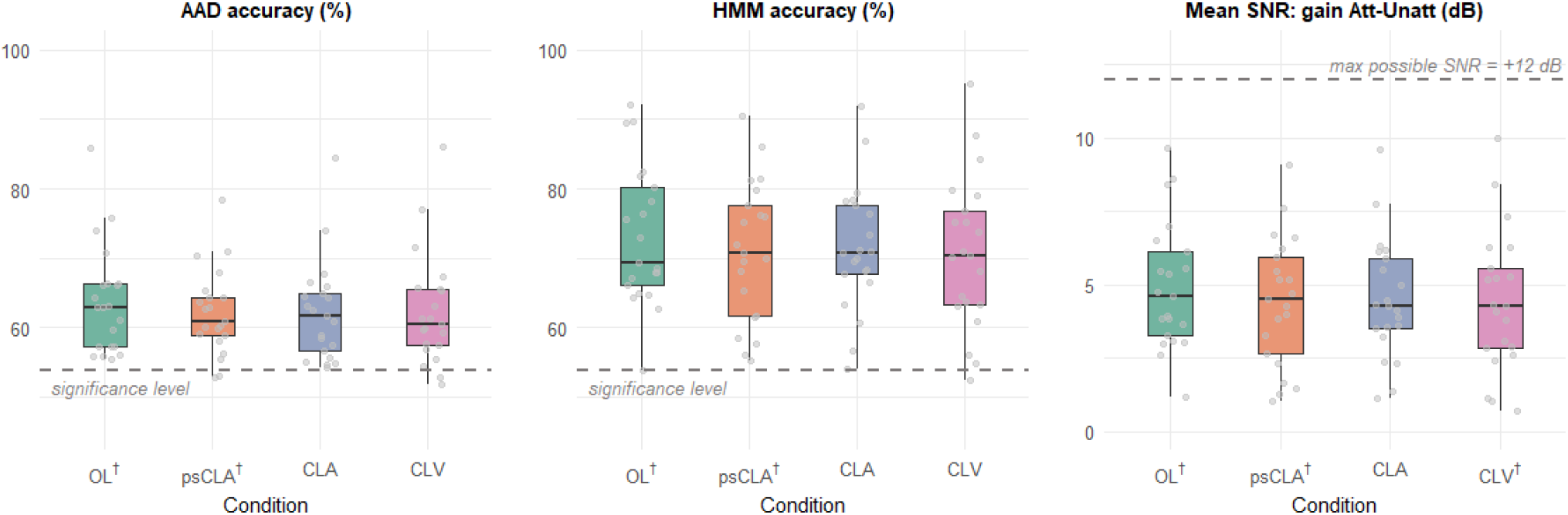
Performance metrics obtained with the online sliding window (wl = 5 s, overlap = 4 s) in CLA and CLV. In OL and psCLA, no actual online decoding occured, but posthoc performance metrics were computed with the same AAD decoder and HMM hyperparameters used online in the other conditions, in order to emulate online decoding and facilitate comparison across conditions. These posthoc scores are illustrated with the †-symbol. Per boxplot, each datapoint respresents the mean performance score of the respective condition in a single participant. Subjects with online SS and SI decoders are pooled together per condition/boxplot (as no significant effect of decoder type was found).

Notably, the HMM accuracy is consistently higher than the AAD accuracy, which is an inherent property of our step-wise gain control strategy and is in line with the results obtained in Heintz et al^28^. This happens because when the decoder makes errors after a series of consecutive correct decisions, the gain of the target speaker will start to decrease, but will still be higher than the gain of the non-target speaker. This holds true until the gains reach the crossover points, when the gain of the non-target speaker starts to dominate (see fig. 4).

The LME model revealed no significant effects of condition or online decoder type (SS/SI) on either metric (*p >* 0.05). Posthoc pairwise comparisons between conditions, adjusted using Tukey’s method, confirmed the absence of statistically significant differences (*p >* 0.05 for all condition pairs in all metrics). The primary source of variance was the participant-level random intercept, with an estimated standard deviation of *σ*_AAD__accuracy_ = 6.95, *σ*_HMM__accuracy_ = 9.04, *σ*_mean__SNR_ = 2.04, indicating substantial individual variability in performance.

### 3.2 Offline decoding (posthoc): audio feedback degrades AAD performance

To evaluate the robustness of the findings above, we re-processed the data offline using a more standardized AAD performance analysis setting with non-overlapping windows (of 5 s and 60 s) and two distinct subject-independent (SI) decoders and no HMM post-processing. Firstly, the SI decoder used online for subjects 1-10 (trained on the AAD-KUL dataset) was reapplied offline on *all* subjects, and on decoding windows without overlap. Secondly, we considered an SI decoder trained on the present dataset, specifically on all the OL trials, but in a leave-one-subject-out fashion (further called the SI AAD-NFB decoder). Figure 7 illustrates the offline AAD accuracies per decoder and window length.

**Figure 7.**
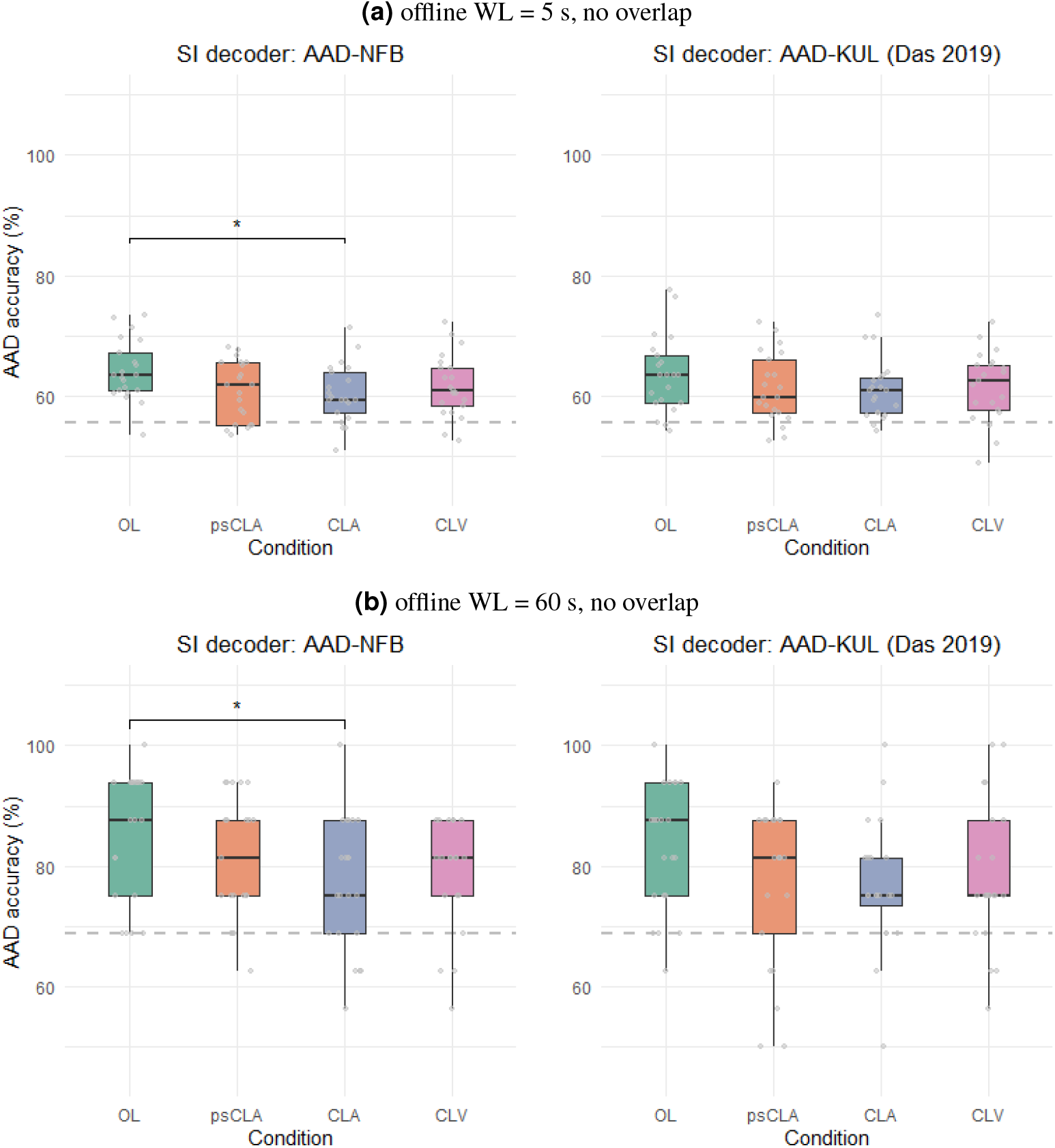
Offline decoding results obtained posthoc with two subject-independent (SI) decoders, after segmenting the data in non-overlapping windows of **(a)** 5 s and **(b)** 60 s. Each datapoint respresents the mean AAD accuracy of the respective condition in a single participant. LME models with posthoc comparisons correction were used to compare scores across all distinct pairs of conditions. Only the statistically significant pairs in the posthoc testing are annotated (* denotes *p*_*adj*_ ∈ [0.01, 0.05), after adjusting for multiple comparisons). All other pairs are not significant. The gray dashed lines indicate the significance levels corresponding to the tested window lengths.

For the offline SI AAD-NFB decoder, the type III ANOVA applied on the fitted LME model revealed a significant main effect of condition both for wl = 5*s* (*F*(3, 59.78) = 3.54, *p* = 0.02) and for wl = 60*s* (*F*(3, 79) = 2.94, *p* = 0.04). Posthoc pairwise comparisons (Tukey-adjusted) showed that the OL condition yielded significantly higher accuracy than the CLA condition (*β* = 3.77, *p* = 0.021 for wl = 5*s* and *β* = 8.63, *p* = 0.03 for wl = 60*s*). Notably, the performance gap between OL and CLA widened as the window length increased from 5 s to 60 s, suggesting that longer integration windows may amplify the observed differences between OL and CLA. The effect of the decoder type (SI/SS) used online for producing neurofeedback was non-significant (*p >* 0.05) for either window length.

In contrast, using the SI AAD-KUL decoder trained on the Das et al. 2019 dataset^26^, no main effect of condition was found (*p* = 0.13 for both window lengths). For wl = 60*s*, the initial LME model did show a significant degradation of AAD accuracy in the CLA condition compared to OL (*β* = −7.74, *p*_*unadj*_ = 0.026), which is also visually detectable (see fig. 7b, right). However, this significant effect did not survive posthoc testing with multiple comparisons correction (*p*_*adj*_ = 0.11). This suggests that while a trend towards lower performance in CLA exists, the high inter-subject variability (shown by the individual data points) likely masked potential differences. The effect of the decoder type used online (SI/SS) was also non-significant (*p >* 0.05) for either window length.

### 3.3 Behavioral results: audio feedback increases both distraction and switching effort

Figure 8 illustrates the mean subjective ratings reported by participants per condition. LME and type III ANOVA modeling revealed a significant main effect of condition only for the scores of ‘Degree of distraction’ (*F*(3, 61.44) = 4.18, *p* = 0.01) and ‘Attention Switching Effort’ (*F*(3, 61.12) = 4.88, *p* = 0.004), suggesting that the experimental conditions imposed distinct cognitive demands on the participants. Posthoc pairwise comparisons (Tukey-adjusted) revealed that the CLA condition was perceived as significantly more distracting than both the OL (*p* = 0.01) and psCLA (*p* = 0.02) conditions. Furthermore, participants reported a significantly higher effort to switch attention in CLA compared to OL (*p* = 0.004) and CLV (*p* = 0.02). While not reaching statistical significance in posthoc contrasts (*p*_*adj*_ *>* 0.05), CLA also trended toward the lowest speech intelligibility (*β* = −0.5, *p*_*unadj*_ = 0.03) and the highest listening effort (*β* = 0.54, *p*_*unadj*_ = 0.04) relative to the baseline OL.

**Figure 8.**
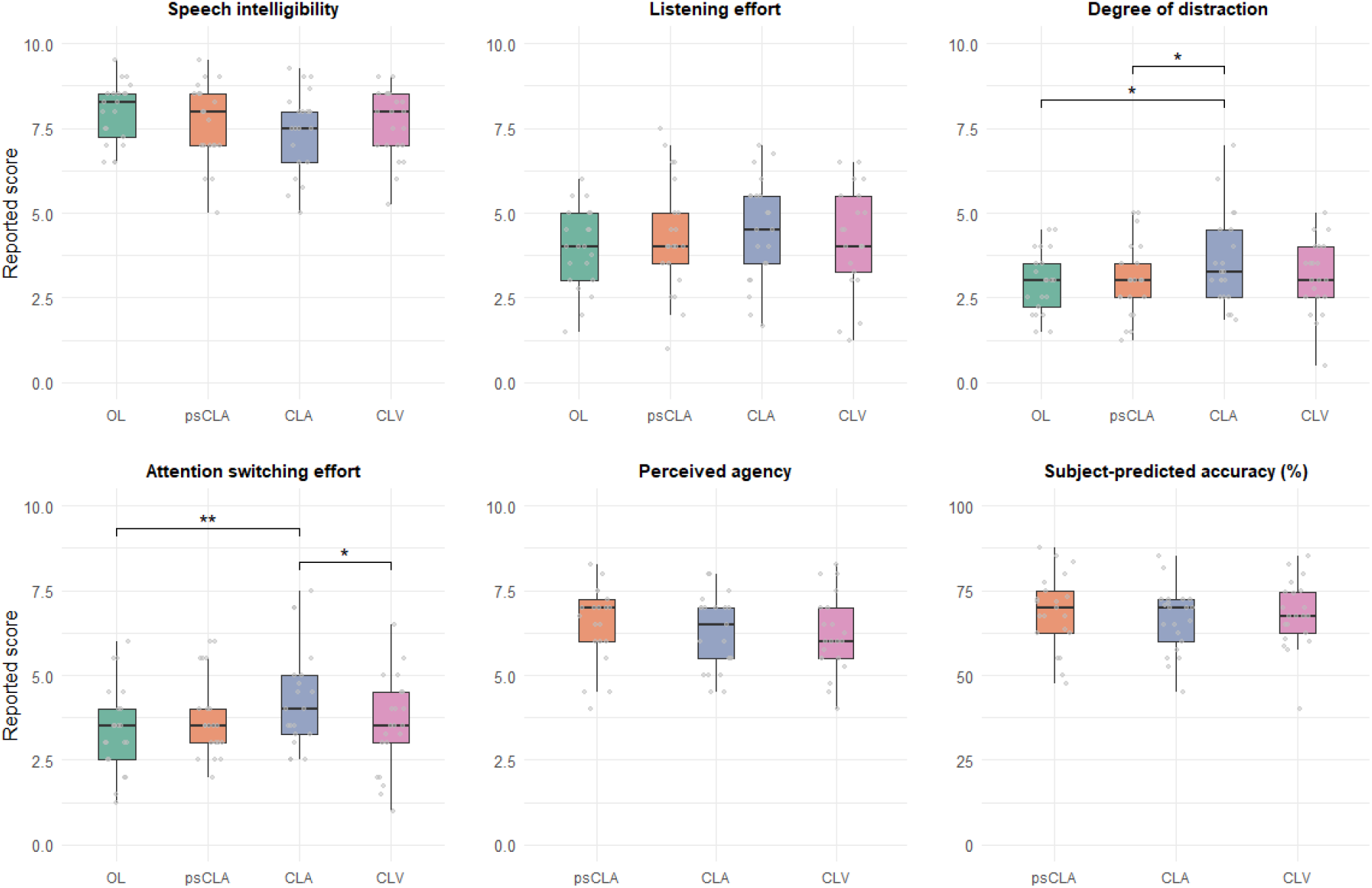
Behavioral results per condition. Each datapoint represents the per-subject reported scores averaged across the two trials of the respective condition. We used LME models with posthoc pairwise comparisons to compare scores per each distinct pair of conditions. The statistically significant pairs are annotated with (**) for *p*_*adj*_ ∈ [0.001, 0.01) and with (*) for *p*_*adj*_ ∈ [0.01, 0.05). All other pairs were not statistically significant.

Altogether, these results suggest that the transition between OL and CLA significantly altered the perceived difficulty of maintaining focus and shifting attention, with CLA typically rated as more demanding than OL.

These results inform us about the subjective experience of the user, with the CLA condition appearing to be the most difficult in terms of ability to stay focused and switch the auditory attention. We suspect that switching attention in CLA is particularly effortful because during the switch, subjects are not only faced with a change of context and speaker particularities (e.g., voice, rhythm, intonation), but also a change in the level of audibility. In particular, the AAD decoding and gain control have an inherent algorithmic delay, which means that it takes some time before the gain of the new target speaker transitions from minimum to maximum amplification (in the best case scenario) after the switching cue is shown. Notably, switching attention in CLV is not significantly harder than in OL (*p >* 0.05), thus confirming that the feedback modality can differentially impact the attention switching performance (in CLV, the visual feedback does not interfere with the switching ability in the auditory domain as both speakers are presented with the same volume).

Interestingly, despite the subjective increase in distraction and attention switching effort in the CLA trials, participants reported similar scores of ‘Perceived agency’ and ‘Predicted accuracy’ in all feedback conditions (*p >* 0.05). This indicates that subjects perceived all feedback conditions in a similar manner, thus reinforcing the ‘perceived’ authenticity of the psCLA condition.

### 3.4 Objective and subjective metrics of performance are correlated

One of our research questions is to what extent the objective performance measures predict the subjective experience of the user when operating the system. Thus, we performed a linear regression analysis to evaluate the relationship between the online/posthoc HMM accuracy (at WL= 5*s*, overlap= 4*s*) and the behavioral scores of ‘Predicted accuracy’ and ‘Perceived agency’ across trials and subjects within each neurofeedback condition (CLV, CLA, psCLA), with the results illustrated in fig. 9.

**Figure 9.**
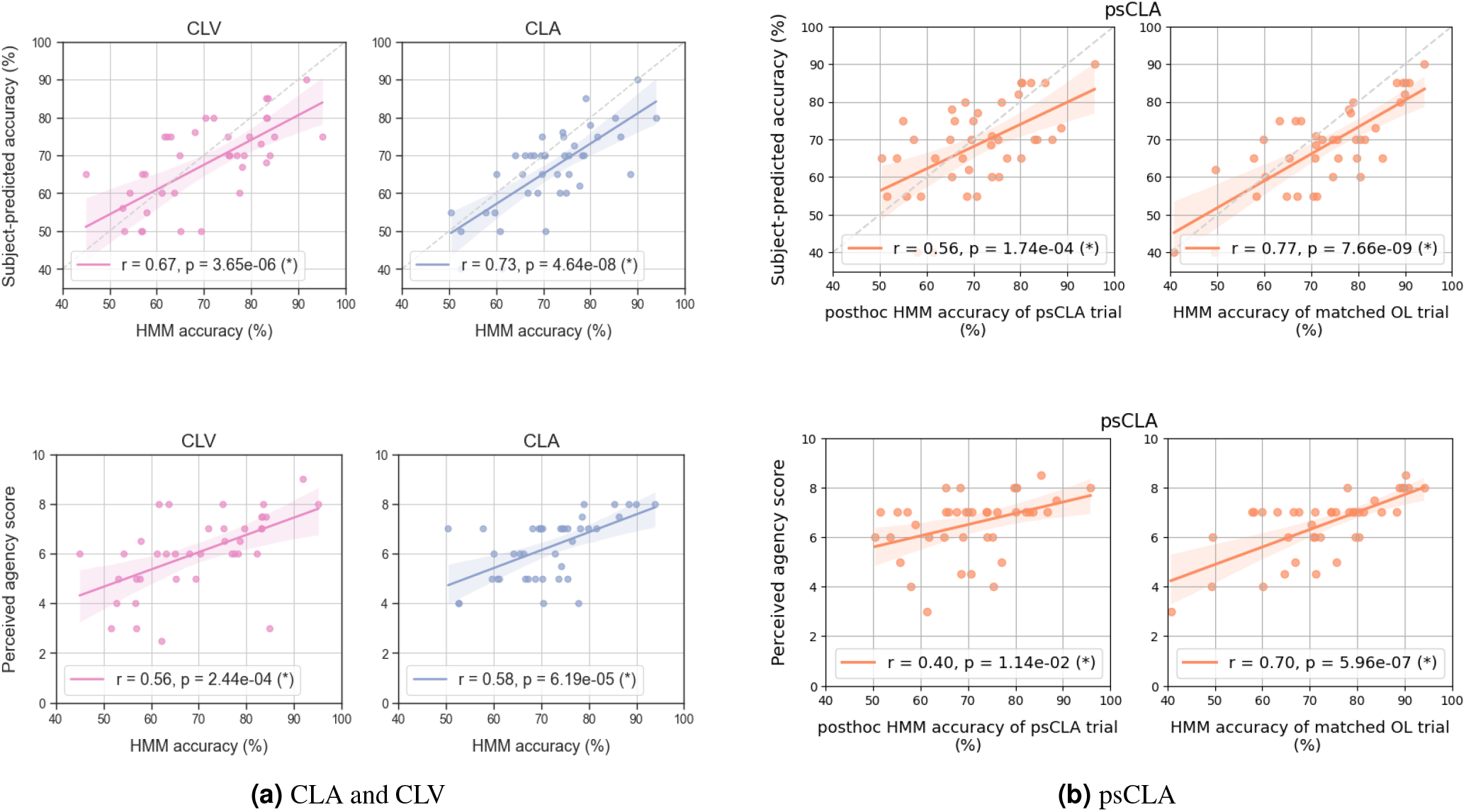
Positive associations between objective metrics (online/posthoc HMM accuracy) and behavioral metrics (‘Subject-predicted accuracy’ and ‘Perceived agency’ score) in **(a)** the online feedback conditions CLA and CLV and **(b)** the pseudo-feedback condition, i.e., psCLA. For the latter, two objective metrics were considered: the HMM accuracy calculated *posthoc* on the data recorded from each psCLA trial and the HMM accuracy of the on the data recorded from the matched OL trial, whose gain trajectory was actually presented “online” (unbeknownst to the participants). Per condition, a linear regression model was fitted to correlate the two variables. Each datapoint represents the respective metrics for a single trial and subject. The gray dashed diagonal line in the top plots denotes the boundary for which the objective HMM accuracy coincides with the subjective predicted accuracy.

The models indicated significant positive linear relationships (*p <* 0.05) between the HMM accuracy and the behavioral scores (the higher the HMM accuracy, the higher the ‘Predicted accuracies’ and ‘Perceived agency’ scores), suggesting that subjects are able to estimate their general performance in all feedback trials and that they feel more in control when the decoding is objectively more accurate.

In particular, in psCLA, positive correlations are also observed between the behavioral scores and the HMM accuracy calculated posthoc (fig. 9b, left column), yet with lower correlation values than in CLV and CLA. However, as expected, stronger associations are observed between the behavioral scores in psCLA and the accuracies of the matched OL trials (fig. 9b, right column), whose underlying gain trajectories were actually presented and perceived by the subjects in psCLA.

Interestingly, the top row of fig. 9 reveals a slight but consistent trend of subjects underestimating their performance across all feedback conditions: the majority of the datapoints concentrate below the first diagonal (where predicted accuracy equals actual accuracy), suggesting that users are slightly more critical of their own performance than the AAD-HMM system. This could be because they misinterpret, based on the perceived feedback, that they make more mistakes than they actually

Here, we chose to focus on the HMM accuracy and not on the AAD accuracy, since the feedback that the subjects observe is determined by the HMM outputs. Thus, the HMM accuracy metric more closely reflects the participant’s subjective experience. We note that similar positive and significant associations were observed between the behavioral scores and AAD accuracies, albeit with lower Pearson correlation values (results omitted). Overall, this analysis informs us that better decoders (with higher decoding accuracy) are required for a more positive user experience, which essentially translates to a high sense of control and an enhanced ability to predict the system performance.

### 3.5 Impact of acoustic gains on AAD performance

To disentangle the effects of neurofeedback-driven neural modulation from the potential influence of acoustic gain on AAD performance, we evaluated the relationship between the presented attended gain 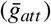 and decoding accuracy.

Figure 10 (a) illustrates the grand-average distribution (over all subjects) of the attended speaker gains presented in the psCLA trials. For *WL*_*gain*_ = 1 s, the histogram distribution directly reflects the online gain updates from the experimental trials. As *WL*_*gain*_ increases, the distributions represent the average gain over longer intervals, resulting in a lower total count of windows but maintaining a consistent trend: the majority of windows fall within the highest gain bin of [2, 6] dB (note that the y-scaling across window lengths is not uniform, to enhance the bins visibility relative to each other). This confirms that even in the decoupled psCLA condition, the subjects were presented with a predominantly enhanced target speaker, thus preserving the apparent authenticity of the neurofeedback task (as intended).

**Figure 10.**
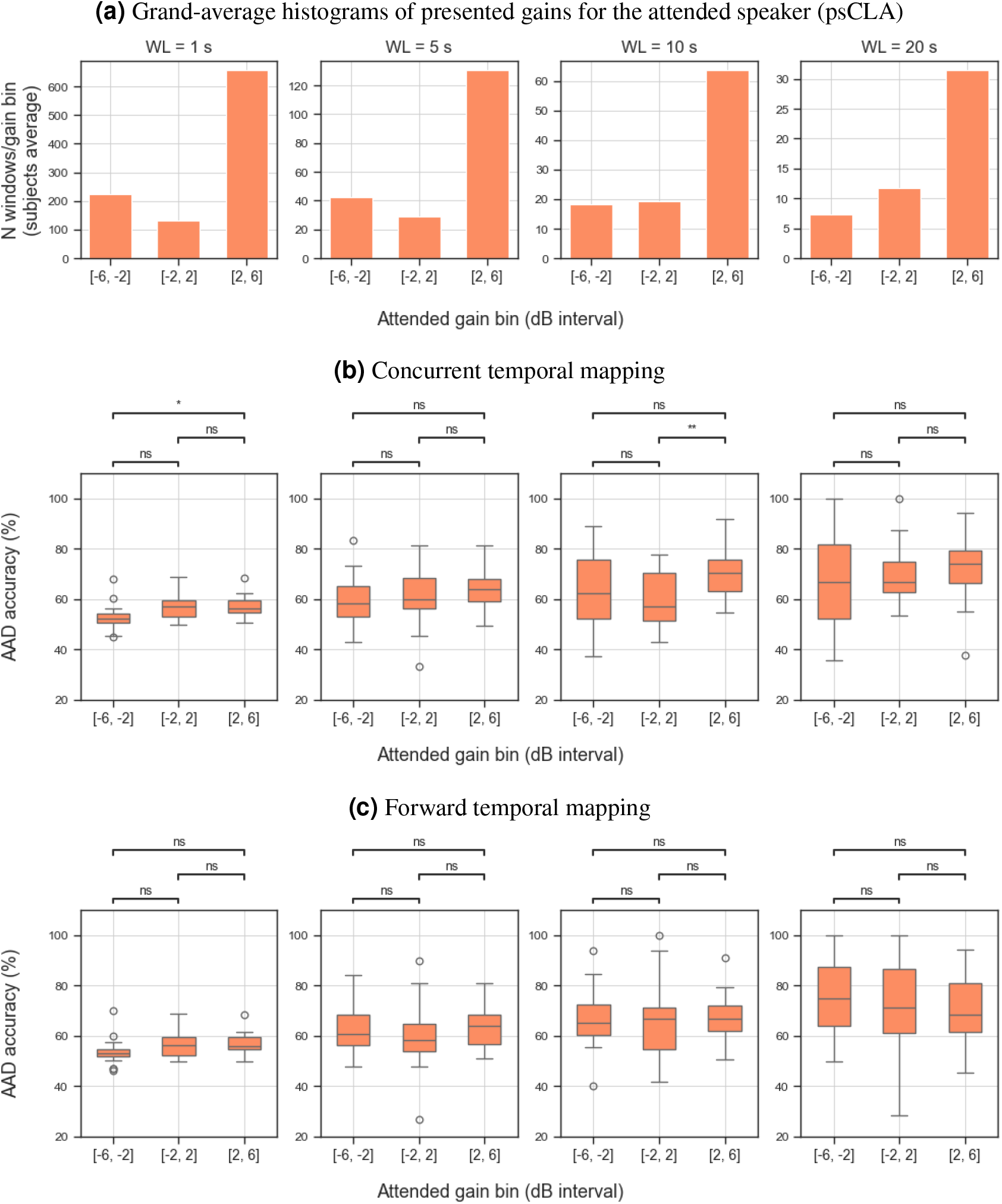
Temporal mapping analysis (posthoc) that probes for acoustic gain effects on the AAD accuracy (unidirectional causality) in the psCLA trials. **(a)**: the grand-average histograms (over subjects) of the online presented gains averaged across the various analysed window lengths (WL). **(b) and (c)**: the AAD accuracy scores per gain bin for the concurrent and forward temporal mapping, respectively, with each boxplot aggregating datapoints across subjects. Each column corresponds to the WL marked on the top row. Per WL and temporal mapping, Wilcoxon signed-rank tests with multiple comparison correction is used to compare accuracy scores between pairs of gain bins. Statistical significance is annotated with (*) for *p* ∈ [0.01, 0.05) and (**) for *p* ∈ [0.001, 0.01); n.s. = not significant

The results of the temporal mapping analyses probing both instantaneous (*ℳ*_1_) and delayed (*ℳ*_2_) effects of acoustic gains on AAD accuracy are summarized in fig. 10(b) and (c).

For the concurrent temporal mapping (*ℳ*_1_), there appears to be no consistent trend of higher gains leading to higher AAD accuracies. While a statistically significant difference emerged between the lowest and highest gain bins at *WL*_*gain*_ = 1 s, and between the middle and highest bins at *WL*_*gain*_ = 10 s, these effects were sporadic and did not persist across all window lengths. Given the high variance and the degraded SNR of the AAD correlations values, especially in the shorter windows (1 s), these isolated significant differences likely represent stochastic fluctuations rather than a robust psychophysical effect.

The forward mapping (*ℳ*_2_) analysis revealed no significant influence of the current gain on the AAD accuracy of subsequent windows across any tested *WL*_*gain*_. The AAD accuracies are statistically on par across all three gain bins (*p >* 0.05), suggesting that acoustic enhancement or suppression does not directly influence the neural tracking of the target speaker in the immediate future. As *WL*_*gain*_ increases, this effect is expected, since the time-difference between the windows becomes too large for any tangible influence of acoustic gains on future neural activity.

## 4 Discussion

This study proposed a novel AAD protocol with online neurofeedback presented in various sensory modalities to probe for short-term learning effects on objective and subjective performance in a selective auditory attention task. Besides the technical validation, we performed both a quantitative and qualitative evaluation of the system through EEG-based decoding performance and behavioral ratings.

Within the closed-loop AAD research landscape^7,8,22,23^, our study is the first to date to include experimental conditions with various feedback modalities in order to evaluate the potential modality-specific neurofeedback benefit w.r.t. to the reference open-loop condition and to disentangle effects of genuine and pseudo-feedback presentation. Furthermore, we here focus on the specific research question of whether an objective improvement in attention decoding can happen as a result of a short-term compensatory neural adjustment to correct for the decoder’s errors, which is what we define as the “neurofeedback benefit”. Our AAD protocol was also realistic, including attention switches with different occurence frequencies.

One surprising finding is that, contrary to our expectations, the type of the online decoder (SS/SI) did not significantly impact the online/offline performance metrics, nor the subjective behavioral outcomes. This was further supported by a secondary LME analysis, which revealed no significant interaction effect between the experimental condition and the decoder type (*p >* 0.05) for any online performance metric (AAD accuracy, HMM accuracy, or mean SNR). These results suggest that the SI decoder, when trained on a sufficiently large and diverse corpus like AAD-KUL^26^, can capture robust, generalized neural signatures of auditory attention that are, in this specific case, functionally equivalent to those identified by the SS decoders.

Nevertheless, perhaps the most striking finding in this study is the significant decrease in decoding accuracy during the CLA condition compared to the OL baseline, although this effect was only observed in an offline posthoc analysis with other decoding settings than those used to generate the feedback. Although our online results are in line with prior results from Aroudi et al.^7^, where no performance difference was observed between OL and CL conditions, our offline results are in opposition with both Aroudi et al.^7^ and our initial hypothesis (stating that acoustic neurofeedback could be beneficial for enhancing attention to the target speaker and its decoding). Below we speculate on the possible reasons for the observed inverse effect.

### 4.1 The behavioral cost of auditory feedback

While neurofeedback is intended to reinforce the neural tracking of the target speaker by maximizing its volume, our results suggest that continuous auditory feedback may introduce a modality-specific cognitive load that interferes with the primary task, acting as a “distractor”^21^.

Overall, the behavioral results (from fig. 8) directly corroborate with the objective decoding results obtained posthoc (from fig. 7): the degree of distraction and switching effort were highest in CLA, while the objective metric of AAD accuracy was lowest. There was also a slight tendency of higher listening effort reported in CLA, although not statistically significant. One possible explanation lies in the dual-task nature of the CLA condition: participants must maintain focus on the target speaker while simultaneously monitoring the fluctuating volume (gain) of that speaker as a proxy for their own performance. In an attention paradigm where the audio signal is crucial for performing the main task, the changing acoustic gains add an extra layer of complexity, which becomes very taxing in the cases of wrongly decoded attention and during switch periods, when negative SNRs are reached. In such cases, the participant may have to exert extra cognitive and attentional effort to maintain focus on the acoustically-suppressed target speaker. Although the average online SNR was positive in the CLA condition (see fig. 6), its potential benefit may be offset by confusion and increased effort when the wrong speaker is amplified. Even during short trials, the constant awareness of momentary lapses in decoding and their effect on the target speaker gains may negatively impact the participants’ motivation and ability to pay attention, leading to an overall degradation in attention decoding performance.

Furthermore, the decoding and behavioral performance in the CLV condition did not significantly differ from the baseline OL, which suggests that shifting the feedback to the visual modality is less distracting and likely bypasses the bottleneck of the limited auditory processing abilities in the dual-task auditory paradigm. In light of these results, we posit that online visual feedback does not introduce additional confounds w.r.t. the primary auditory task, thus allowing participants to be informed about their performance without affecting the neural representation of the acoustic target.

### 4.2 High decoder quality as a prerequisite for neurofeedback learning effects

We suspect the quality of the trained AAD decoders and their accuracy to play a major role in online paradigms with neurofeedback and that a minimal threshold of performance is required to facilitate learning and performance improvement through behavioral/neural adaptation. A supplementary analysis of the correlation between the AAD accuracies in baseline OL vs. CLA indeed confirms there is a positive association (statistically significant) between the two performance metrics (*r* = 0.8, *p* = 1.36*e* − 5 for the online results with the respective SS/SI decoders and *r* = 0.68, *p* = 7.68*e* − 4 for the offline SI decoder trained on the AAD-KUL dataset^26^), suggesting that better baseline performance is associated with improved performance in a closed-loop condition.

In light of our results, we suspect that our decoders did not reach a sufficient baseline performance to elicit sufficiently strong positive NFB effects in either feedback modality. A decoder that makes the subjects face wrong predictions frequently can lead to a negative self-reinforcing effect: inaccurate feedback may induce the subjects to feel they lose control, which in turn distracts or demotivates them, leading to a further drop in decoding performance. Thus, the use of suboptimal decoders brings the risk that the system is working against the user, which is the exact opposite to the intended outcome.

It is also plausible that mere behavioral compensation and enhanced attention levels might not be sufficient to bypass a fixed and inherently suboptimal decoder. In fact, the success of a closed-loop system relies on the synergy between two agents: the user and the decoder. In the ideal case, both would adapt over time to optimize performance, which is known as ‘co-adaptation’^29^. Future work could replace the fixed decoders with time-adaptive decoders (which are retrained online with the latest available EEG data^30,31^) and evaluate their potential performance gains in an online neurofeedback AAD paradigm. Finally, we note there is a slight discrepancy between the online and offline decoding results: while online performance was consistent across conditions, offline analysis revealed a significant drop in CLA accuracy. We attribute this to two complementary mechanisms.

First, the online system utilized a sliding-window approach and HMM-based smoothing. The HMM likely buffered the system against the attentional fluctuations induced by the auditory feedback, maintaining equally high accuracy online despite a degraded neural signal in CLA compared to the OL trials. When the data were re-analyzed offline without this temporal smoothing, the penalizing cost of auditory feedback became apparent regardless of the decision window length. In addition, with HMM postprocessing applied offline on the non-overlapping windows, LME modelling and posthoc tests showed no significant differences in the HMM accuracy metric across conditions for either window length (5 vs. 60 s) - results omitted.

At the same time, the mismatch between online and offline decoder settings could provide an alternative explanation. This mismatch implies that the decoding errors of the online and offline decoders do not align in time, hence the errors made by the offline decoder will not (always) be compensated for by the participants (no active error-correction happens offline), which could explain the lower accuracy in the offline decoding setting compared to the online neurofeedback setting. Thus, we suspect that a *negative* feedback loop likely exists in the online CLA condition, which stabilizes the interaction between the decoder and the user and explains why the online results in CLA are on par with easier conditions: when the decoder begins to fail, the resulting change in audio gain alerts the participant, who then exerts extra cognitive effort to re-focus and “correct” the system. This active error correction thus appears to stabilize the online performance in CLA.

### 4.3 Limitations of our neurofeedback protocol

Other reasons for the lack of positive neurofeedback effects could be related to the nature of our protocol. In particular, our single-session approach might be too long and tiring for subjects to learn how to optimally operate the closed-loop system and deal with its prediction errors. Neurofeedback effects typically occur following training paradigms that spread over multiple days, weeks or months^32,33^, while ensuring shorter per-session durations, to avoid exhaustion. Furthermore, the heterogeneity of the presented conditions and the block design with interleaved audio and visual feedback trials might have confused the participants and force them to use different mental strategies, adding to the overall task complexity.

### 4.4 The pseudo-feedback was perceived as genuine

The correlation analysis between objective and subjective performance metrics revealed positive significant relationships across all feedback conditions (cf. section 3.4). In the genuine NFB conditions (CLA and CLV), this alignment is expected, as the frequent fluctuations of auditory gain and visual slider position provide immediate cues regarding decoding success.

Perhaps more surprising was that the same positive association, although to a lower extent, holds true for the psCLA condition, given that the presented feedback was actually disconnected from the subjects neural signals. In addition, LME statistical models with posthoc constrasts revealed no significant difference in the reported score of ‘Perceived agency’ between the CLA and psCLA condition. Altogether, this indicates that the subjects perceived the psCLA condition as a genuine CLA condition.

Although counterintuitive, this can be explained by the fact that the feedback shown in psCLA was designed to be realistic, by using gain trajectories from an OL trial. Importantly, in our experimental design, we explicitly enforced that both the OL and psCLA trial have matching attention and switching patterns. As a sanity check, an additional regression analysis (depicted in supplementary **??**) confirms that the AAD/HMM accuracy of the psCLA trial (computed posthoc) correlates positively and significantly with the AAD/HMM accuracy in the matched OL trial. Although this correlation doesn’t necessarily imply causality, the two trials (OL and psCLA) do share the same underlying decoder and attention switching pattern within participant. Thus, it is plausible that a good decoder which yields a stable gain trajectory and a higher accuracy in the OL trial will also enable a higher accuracy in the psCLA trial (in which the OL gain trajectory was presented online), thus creating the illusion of feeling more in control.

### 4.5 Decoding performance is robust to acoustic gain fluctuations

Our causal analysis on the effect of presented gains on the AAD accuracies did not reveal a consistent significant effect across window lengths and temporal mappings. Collectively, these findings suggest that within the gain range tested ([−6, 6] dB), the AAD accuracy is robust to fluctuations in the acoustic SNR between the target and the masker speakers. However, we highlight one caveat of this causal analysis - the unbalanced distribution of decision windows across gain bins, which is a direct artefact of our acoustic presentation with quickly-varying gains. Critically, the high density of the rightmost gain bins relative to the others (see fig. 10) introduces a bias in the statistical power that may affect the robustness of our accuracy estimates across gain bins. Future AAD protocols should consider sufficiently long trials with an even distribution of SNRs between the target and the masker speaker (ideally with fixed SNRs per trial, to avoid transitory neural responses to changing SNRs) in order to confirm these observed effects.

Despite these limitations, our results corroborate with the findings reported by Ding and Simon (2012)^1^, who showed that the neural representation of the attended speaker is stable and invariant to the loudness of both the attended speech stream and the background speech. Moreover, Verschueren et al.^34^ also found that neural envelope tracking (quantified as the correlation between the reconstructed envelope and the acoustic envelope of the attended speech) is generally robust to changes in stimulus intensity, provided the decoder is trained on the same intensity level as the test stimulus.

In light of these findings, we interpret the absence of a systematic gain–accuracy relation in our study as evidence that the decoder’s performance is not driven by the bottom-up acoustic prominence of the attended speaker, but most likely by the subject’s top-down attentional state^35,36^. Moreover, if we extrapolated this interpretation to the CLA condition, we could attribute the observed performance decrements in CLA to the feedback-induced cognitive load rather than the acoustic SNR manipulations.

### 4.6 Conclusion

This study investigated whether neurofeedback (NFB) can improve single-session attention decoding performance and user experience in a realistic two-speaker AAD paradigm with attention switches. Nineteen participants performed a selective listening task in four conditions: open-loop without feedback (OL), closed-loop with auditory feedback (CLA), closed-loop with visual feedback (CLV), and closed-loop with pseudo-auditory feedback (psCLA) using precomputed gain trajectories from a matched OL trial. Online analysis with 5 s sliding windows and HMM smoothing showed comparable decoding performance across conditions. In contrast, offline posthoc decoding with two different subject-independent decoders (no HMM applied) and non-overlapping windows revealed that AAD accuracy in CLA was significantly lower than in OL. Subjective ratings indicated that CLA was perceived as significantly more distracting and required significantly higher switching effort than OL. An additional posthoc offline analysis restricted to the psCLA condition revealed no robust causal effect of presented audio gains on accuracy, suggesting that the reduced performance is not caused by the suppression of the attended speaker due to previous decoding errors. Altogether, these results demonstrate that, within the tested single-session paradigm with rapidly varying feedback cues, auditory neurofeedback does not enhance and can even degrade AAD performance, while increasing subjective distraction and switching effort. This further indicates that suboptimal auditory feedback can impede learning and risks working against the user, increasing the cognitive load instead of facilitating attention decoding. We propose that more accurate and stable decoders, as well as longitudinal, multi-session training protocols may be critical prerequisites for observing beneficial neurofeedback effects in closed-loop AAD.

## Data availability

The EEG dataset generated and analyzed during the current study is publicly available in the following Zenodo repository: [LINK WILL BE AVAILABLE UPON ACCEPTANCE].

## Author contributions statement

I.R., S.G., A.B., and T.F. conceptualized the study. N.H. and A.B. developed the real-time gain-control algorithm. I.R. performed the technical implementation, data collection, formal data analysis and drafted the initial manuscript. All authors contributed to general discussions, subsequent manuscript revisions and approved the final version.

## Additional information

## Acknowledgements

We are thankful to Ann Agterberg for assisting with participants recruitment and data collection and to Abdulahad Kancan for technical assistance with software development.

Financial support was provided by the Research Foundation Flanders (SBO mandate 1S14922N for I. Rotaru, SBO mandate 1S31522N for N. Heintz, junior posdoctoral mandate 1242524N for S. Geirnaert and FWO projects G081722N, G026026N), the Flemish Government (AI Research Program), Internal Funds KU Leuven (projects C3/25/017, IDN/23/006 and C14/25/108), and the European Research Council (ERC) under the European Union’s research and innovation programme (grant agreement No 101138304). Views and opinions expressed are however those of the author(s) only and do not necessarily reflect those of the European Union or any of the granting authorities. Neither the European Union nor the granting authorities can be held responsible for them.

