## Supplementary Information for "When feedback backfires: investigating neurofeedback effects in a closed-loop auditory attention decoding paradigm"

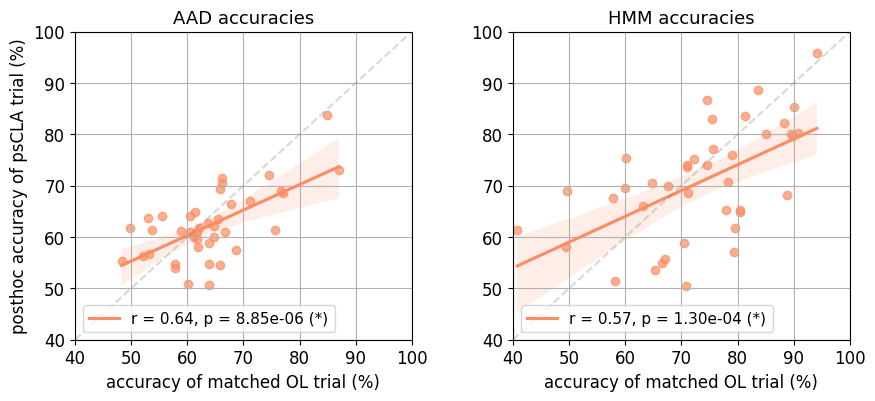
**Supplementary Fig. 11**
Positive association between OL and psCLA decoding performance. Each datapoint represents the respective metrics for a single trial and subject.

**List of behavioral questions presented in the Neurofeedback-AAD study**

| **Subjective attribute** | **Behavioral question (English translation)** | **Reference values** | **Trials where asked** |
| --- | --- | --- | --- |
| Stimulus comprehension | Content-related | - | All trials: OL, CLA, psCLA, CLV |
| Speech intelligibility | Overall, how much of the speech produced by the green target speaker(s) did you hear and understand in total?   Please answer with a number between 0-10. | 0: I did not hear and understand any speech produced by the target speaker(s)  5: I heard and understood about half of the speech produced by the target speaker(s)  10: I heard and understood all the speech that was produced by the target speaker(s) | All trials |
| Listening effort | How much effort did it take you to pay attention and listen to the green speakers and ignore the black speaker(s)?  Please answer with a number between 0-10. | 0: low effort (it was very easy to attend the green and ignore the black speakers)  5: medium effort (it was somewhat difficult to attend the green and ignore the black speakers)  10: high effort (it was very difficult to attend the green and ignore the black speakers) | All trials |
| Degree of distraction | Overall, how much time of the entire trial were you distracted by the black speaker(s)?  Please answer with a number between 0-10. | 0: during the trial, I was never distracted by the black speaker  5: during the trial, I was distracted by the black speaker about 50% of the time  10: during the trial I was always distracted by the black speaker | All trials |
| Switching effort | How much effort did it take to switch attention between the 2 speakers?  Please answer with a number between 0-10. | 0: low effort (rather easy to switch)  5: medium effort (somewhat difficult to switch)  10: high effort (very difficult to switch) | All trials |
| Perceived agency (sense of control) | Based on the feedback you received throughout the trial (audio/visual feedback), to what degree did you feel you were able to control the system?  Please answer with a number between 0-10. | 0: I could not control the system at all  5: I could somewhat control it  10: I could fully control it | CLA, CLV, psCLA |
| Perceived accuracy | Based on the audio/visual feedback you received, can you predict the accuracy score for this trial?  Accuracy = percentage of time when the system correctly decoded your auditory attention to the green speaker  Please answer with a percentage between 0-100 %. | 0: the system never correctly predicted my auditory attention  50: the system correctly predicted my auditory attention in about 50% of the trial  100: the system correctly predicted my auditory attention in about 100% of the trial | CLA, CLV, psCLA |
